# Predicting the Outcomes of New Short-Course Regimens for Multi-Drug Resistant Tuberculosis Using Intrahost and Pharmacokinetic-Pharmacodynamic Modelling

**DOI:** 10.1101/369991

**Authors:** Tan N Doan, Pengxing Cao, Theophilus I Emeto, James M McCaw, Emma S McBryde

## Abstract

Short-course regimens for multi-drug resistant tuberculosis (MDR-TB) are urgently needed. Limited data suggest that the new drug, bedaquiline (BDQ), may have the potential to shorten MDR-TB treatment to less than six months when used in conjunction with standard anti-TB drugs. However, the feasibility of BDQ in shortening MDR-TB treatment duration remains to be established. Mathematical modelling provides a platform to investigate different treatment regimens and predict their efficacy. We developed a mathematical model to capture the immune response to TB inside a human host environment. This model was then combined with a pharmacokinetic-pharmacodynamic model to simulate various short-course BDQ-containing regimens. Our modelling suggests that BDQ could reduce MDR-TB treatment duration to just 18 weeks (four months) while still maintaining a very high treatment success rate (100% for daily BDQ for two weeks, or 95% for daily BDQ for one week during the intensive phase). The estimated time to bacterial clearance of these regimens ranges from 27 to 33 days. Our findings provide the justification for empirical evaluation of short-course BDQ-containing regimens. If short-course BDQ-containing regimens are found to improve outcomes then we anticipate clear cost-savings and a subsequent improvement in the efficiency of national TB programs.

## INTRODUCTION

Multi-drug resistant tuberculosis (MDR-TB) is an emerging public health crisis. Globally, around 480,000 people develop MDR-TB and 190,000 people die from the disease each year (1). Currently recommended (conventional) regimens for MDR-TB treatment are complex, long (at least 20 months), expensive, poorly tolerated and have a modest global treatment success rate of 52% (1). In order to improve adherence to anti-TB treatment, the World Health Organization (WHO) promotes the use of directly observed therapy (DOT), in which patients undertake medications under the direct observation of a health care provider (2). However, DOT is difficult and costly to implement in resource-scarce settings where health infrastructure is poor and access to health care is limited (3). The prolonged duration of conventional regimens of MDR-TB makes the implementation of DOT even more challenging. Strategies aimed at reducing the duration of MDR-TB treatment and the frequency of dosing administration while improving efficacy and safety profiles are therefore desirable (4).

The nine-month short-course regimen (also known as the Bangladesh regimen) has been reported to be highly effective with treatment success rates ranging from 84 to 88%, and better tolerated than the conventional regimens (5-7). The Bangladesh regimen consists of a fourth-generation fluoroquinolone (moxifloxacin, levofloxacin or gatifloxacin), ethambutol (EMB), pyrazinamide (PZA) and clofazimine (CFZ) given throughout the treatment period, and supplemented by kanamycin (KNM), prothionamide (PTH) and high-dose isoniazid (INH) in the intensive phase (5-7). In recent years, a number of new therapeutics aimed at tackling MDR-TB have emerged. In particular, there has been increasing interest in assessing the efficacy of short-course regimens containing bedaquiline (BDQ), which is a novel diarylquinoline compound with anti-mycobacterial effects (8). Two phase II clinical trials have reported that the addition of BDQ to the conventional 20-month background regimen resulted in a higher sputum culture conversion rate at week 24 compared to the conventional regimen alone (9-11). In both trials, BDQ was given at a dose of 400 mg daily for two weeks in the intensive phase of treatment followed by 200 mg three times a week for six (9, 10) or 22 weeks (11). Such a success makes the suggestion of a BDQ-containing regimen that is shorter than the nine-month Bangladesh regimen for MDR-TB treatment promising. However, the feasibility of BDQ in shortening MDR-TB treatment duration remains to be established. It may be ethically and practically infeasible to run clinical trials on every possible regimen for all epidemiological contexts. Accordingly, mathematical modelling and *in silico* simulations – as used extensively for drug regimen optimisation for other diseases such as malaria (12, 13) – provide an excellent platform to predict the outcomes of MDR-TB treatment based on alternative short-course regimens. Findings from these modelling studies can then be used to guide future clinical trials.

For a mathematical model to realistically mimic the evolutionary dynamics of *Mycobacterium tuberculosis* (*Mtb*) in a human host environment and correctly estimate drug effects, it is essential to combine the immune response into pharmacokinetic (PK) and pharmacodynamic (PD) models of anti-TB drugs (14). This is because of the important role that the immune response plays in the proliferation and clearance of *Mtb* in the host environment (15), as well as the complex interactions between the immune response and the pathogen that determine pharmacological effects of a drug (14). To the best of our knowledge, there has been no attempt to link PK-PD models to host immune response models to optimise MDR-TB regimens. In this paper, we first develop an intrahost model for the dynamics of *Mtb* in a human host environment. We then combine this model with PK-PD models of anti-TB drugs to simulate various short-course regimens for MDR-TB, including regimens that contain the new drug BDQ, and estimate their anti-mycobacterial effects.

## MATERIALS AND METHODS

### The model

The model consists of four main components: (i) intrahost dynamics of *Mtb*; (ii) innate and adaptive immune response; (iii) PK; and (iv) PD of anti-TB drugs. We briefly describe each of these components, and how they were combined, below. A full presentation of the model structures, mathematical equations and input variables is provided in the Supplemental Material.

### Intrahost dynamics of *Mtb*

We simulated the evolutionary dynamics of *Mtb* during active TB disease. In order to allow for the differences in bacterial growth rates and their roles in soliciting the immune response and infection dynamics, intracellular and extracellular *Mtb* populations were modelled separately. The intracellular bacteria (BI) refer to *Mtb* that reside within infected macrophages (MI); and *Mtb* existing anywhere other than MI are considered to be extracellular bacteria (BE) (15). BI and BE loads may change due to processes such as natural growth and death, BI released from MI due to cell rupture, bacteria phagocytised by macrophages, and exposure to anti-TB drugs (15).

### Immune response

Major elements of the host immune response including macrophages, cytokines (interleukins IL-4, IL-10, IL-12, interferon-gamma [IFN-γ]) and T lymphocytes (CD4^+^ and CD8^+^ T cells), all of which are critical to the time-course of *Mtb* dynamics (15), were incorporated. Our “reference space” for the model was bronchoalveolar lavage (BAL) fluid, reported to be a good predictor of the lung environment (16, 17). All cells and bacterial load were measured in units per millilitre of BAL. A detailed description of the host immune response model is provided in Section S1 of the Supplemental Material. Briefly, we distinguished three types of macrophages: resting macrophages (MR), activated macrophages (MA) and MI. Activation of MR to become MA requires IFN-γ in the presence of *Mtb.* MR become MI when they are invaded by bacteria and are unable to effectively clear these BI (15). Both MR and MA can phagocytise and kill BE and secrete cytokines but MA are much more efficient at each of these processes than MR (15). *Mtb* that reside within MI can only be killed when host macrophages are lysed by effector T cells (15).

### Pharmacokinetics and pharmacodynamics of anti-TB drugs

For each of the included anti-TB drugs, a PK model was developed to simulate the concentration-time profiles in plasma, epithelial lining fluid (ELF, extracellular concentrations) and alveolar cells (AC, intracellular concentrations). These models have been found to best describe PK data of the respective anti-TB drugs in previous studies (14, 18-25). Except for INH of which the PK model structure included both ELF and AC compartments (14), ELF-to-plasma and AC-to-plasma concentration ratios were used to extrapolate ELF and AC concentrations from simulated plasma concentrations. The PK model structures and variables are described in Section S2 of the Supplemental Material. Between-subject variability was included where available to capture the variability in PK profiles among individuals.

PD models were developed to describe the relationship between *in vivo* drug concentrations (obtained from the PK models) and killing of bacteria. The rate of change of bacterial load over time (*ψ*(*C*)), i.e. the net growth of *Mtb* exposed to an antibiotic concentration (*C*), was implemented by subtracting the kill rate by antibiotics (*μ*(*C*)) from the intrinsic growth rate of the bacterial population in the absence of treatment (*ψ_max_*) (26). A logistic growth function was used to model the intrinsic growth rate of bacteria in the absence of treatment (27). The growth rate takes the following form:

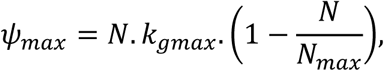

where *N* is the total bacteria count at any given time, *k_gmax_* is the maximal growth rate per unit time, and *N_max_* is the maximum of *Mtb* load.

The antibiotic-mediated kill rate is described by a Hill function with form

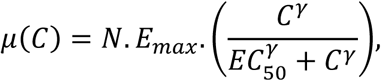

where *E_max_* is the maximal kill rate of each drug per unit time; *EC*_50_ is the antibiotic concentration to achieve half of its maximal kill rate; and *γ* is the Hill coefficient, which is a measure of the steepness of the relationship between the kill rate and drug concentration.

Taken together, the function for the net growth rate of *Mtb* can be written as

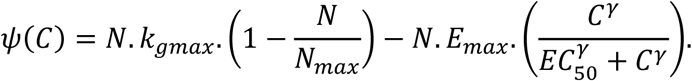

Input values of the PD variables are shown in Section S3 of the Supplemental Material.

### Treatment regimens simulated

We simulated all six standardised regimens with treatment durations ranging from 21 months to nine months (the nine-month regimen is also known as the Bangladesh regimen) that were previously investigated in the landmark clinical trial by the Damien Foundation in Bangladesh (Table1) (6). We then estimated treatment success rates of these regimens, and compared these outcomes with those reported in clinical trials. The treatment success rate was defined as the percentage of the simulated individuals who achieved bacterial clearance during the treatment period and did not exhibit recrudescence within 12 months of treatment completion (6). We defined bacterial clearance as a total bacterial population below the limit of detection of sputum culture method, reported in the literature to be 50 CFU (colony forming unit)/ml (28). Time (in days) to bacterial clearance since treatment commencement was also calculated for patients with successful treatment.

**TABLE 1.** Standardised regimens sequentially used in the treatment of MDR-TB in the clinical trial by the Damien Foundation in Bangladesh (6)

| Regimen | Intensive phase* | Continuation phase 1* | Continuation phase 2* |
| --- | --- | --- | --- |
| 1 | 3 KNM, CFZ, OFX, EMB, INH, PZA, PTH | 12 OFX, EMB, INH, PZA, PTH | 6 EMB, PTH |
| 2 | 3 KNM, CFZ, OFX, EMB, INH, PZA, PTH | 12 OFX, INH, EMB, PZA, PTH |  |
| 3 | 3 KNM, CFZ, OFX, EMB, PZA, PTH | 12 OFX, EMB, PZA, PTH |  |
| 4 | 3 KNM, CFZ, OFX, EMB, INH, PZA, PTH | 12 OFX, INH, EMB, PZA |  |
| 5 | 3 KNM, CFZ, OFX, EMB, INH, PZA, PTH | 12 OFX, INH, EMB, PZA, CFZ |  |
| 6<sup>‡</sup> | 4<sup>§</sup> KNM, CFZ, MXF, EMB, INH, PZA, PTH | 5<sup>§</sup> MXF, EMB, PZA, CFZ |  |

\*Numbers in front indicate months. <sup>‡</sup>Referred to as the “nine-month Bangladesh regimen” in the present study. <sup>§</sup>Moxifloxacin, instead of gatifloxacin, was used in our simulations, due to the lack of pharmacodynamic data for gatifloxacin. CFZ, clofazimine; EMB, ethambutol; INH, isoniazid; KNM, kanamycin; MXF, moxifloxacin; OFX, ofloxacin; PTH, prothionamide; PZA, pyrazinamide.

We also investigated nine-month treatment regimens in which moxifloxacin (MXF) was administered intermittently three times weekly and weekly during the continuation phase. Intermittent dosing of MXF is appealing because MXF is characterised by a long half-life and concentration-dependent antibacterial effects (21, 29). A nine-month regimen in which INH was omitted was also simulated based on recent animal data suggesting that omitting INH may not compromise treatment efficacy (30).

To investigate the potential of BDQ to further shorten MDR-TB treatment to less than nine months, we simulated various short-course BDQ-containing regimens. These regimens consisted of an initial intensive phase with BDQ, MXF, CFZ, PZA, INH and KNM, followed by a continuation phase with BDQ, MXF, CFZ and PZA (Fig. 1). Various durations of treatment (26, 22 and 18 weeks), were investigated and a comparative analysis of their efficacy was undertaken in order to identify highly effective regimens.

**FIG 1.**
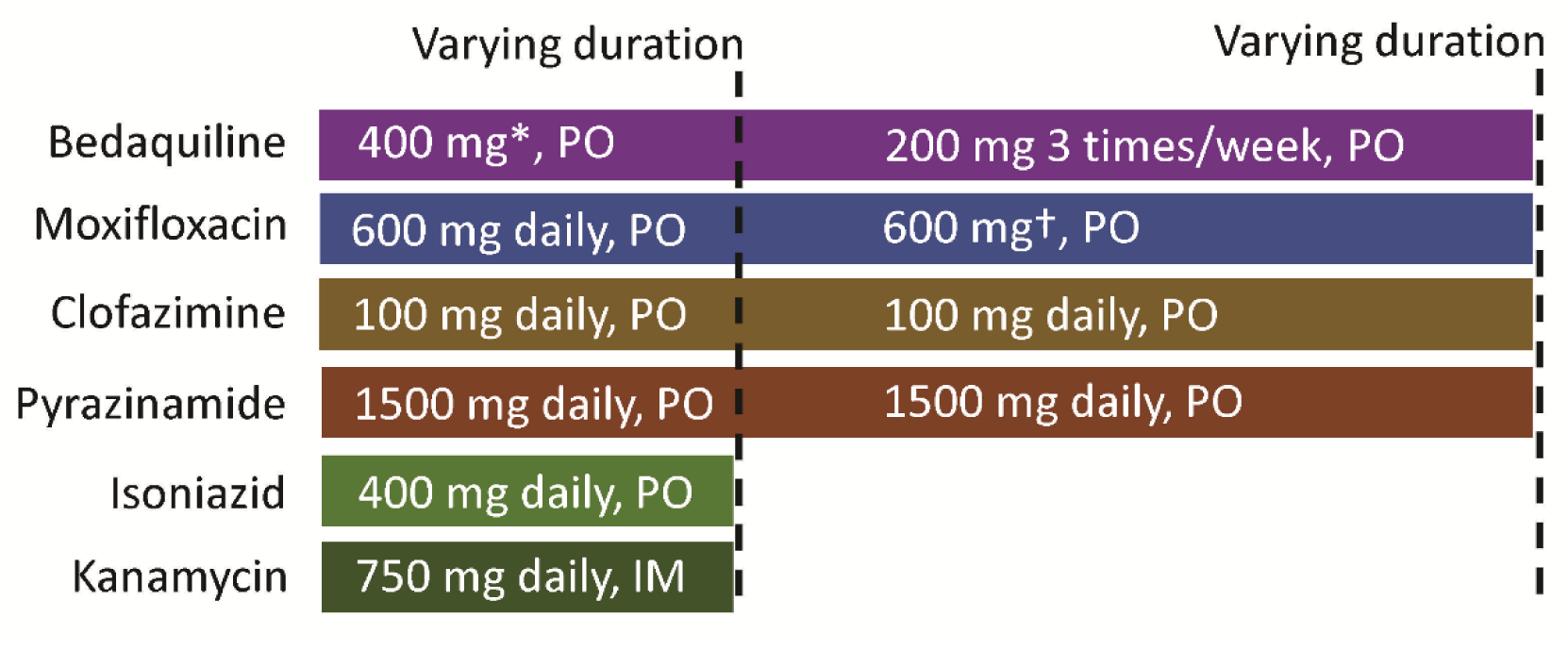
Short-course BDQ-containing regimens. BDQ, bedaquiline; PO, oral administration; IM, intramuscular injection. *BDQ daily and three times weekly during intensive therapy were investigated. †Simulated dosing schedules of MXF included daily, three times weekly or weekly during the continuation phase.

### *In silico* simulation

In order to account for between-subject variability, we simulated a hypothetical cohort of 500 individuals by sampling 500 different parameter sets from parameter space using the Latin Hypercube Sampling method (31). Ranges for the parameters are shown in Tables S1-S3 of the Supplemental Material. For model parameters whose mean values and lower and upper bounds were known, samples were drawn from a modified beta distribution (see Section 4 of the Supplemental Material for a description of the modified beta distribution) to reflect variability in the host population. For non-negative model parameters whose upper bounds were unknown, a log normal distribution was used. Each regimen was evaluated based on the 500 hypothetical patients.

Multivariate sensitivity analysis was performed to investigate the key drivers of bacterial population dynamics and its sensitivity to changes in model inputs. Partial rank correlation coefficients (PRCCs) (32) were calculated to evaluate the strength of the correlation between each outcome measure and each variable. All simulations and analyses were performed using MATLAB (version R2017a, MathWorks, Natick, MA, USA). Ethics approval was not required as this study used data from published literature.

## RESULTS

### Simulation in the absence of treatment

Fig. 2a shows the dynamics of *Mtb* during active TB disease using baseline parameter values, with extracellular *Mtb* representing the majority of the bacteria population. The numbers of extracellular and intracellular bacteria increased and then saturated (i.e. they approached their capacities) on approximately day 150, consistent with the range reported in the literature (150-180 days) (15, 33). The dynamics of BE was primarily driven by the growth rate and the killing by macrophages (Fig. 2b). In contrast, the evolution of BI was mainly governed by the growth rate and the conversion between BE and BI (e.g. invasion of BE into MR and BI released from MI due to MI rupture either naturally or via TB infection) (Fig. 2c).

**FIG 2.**
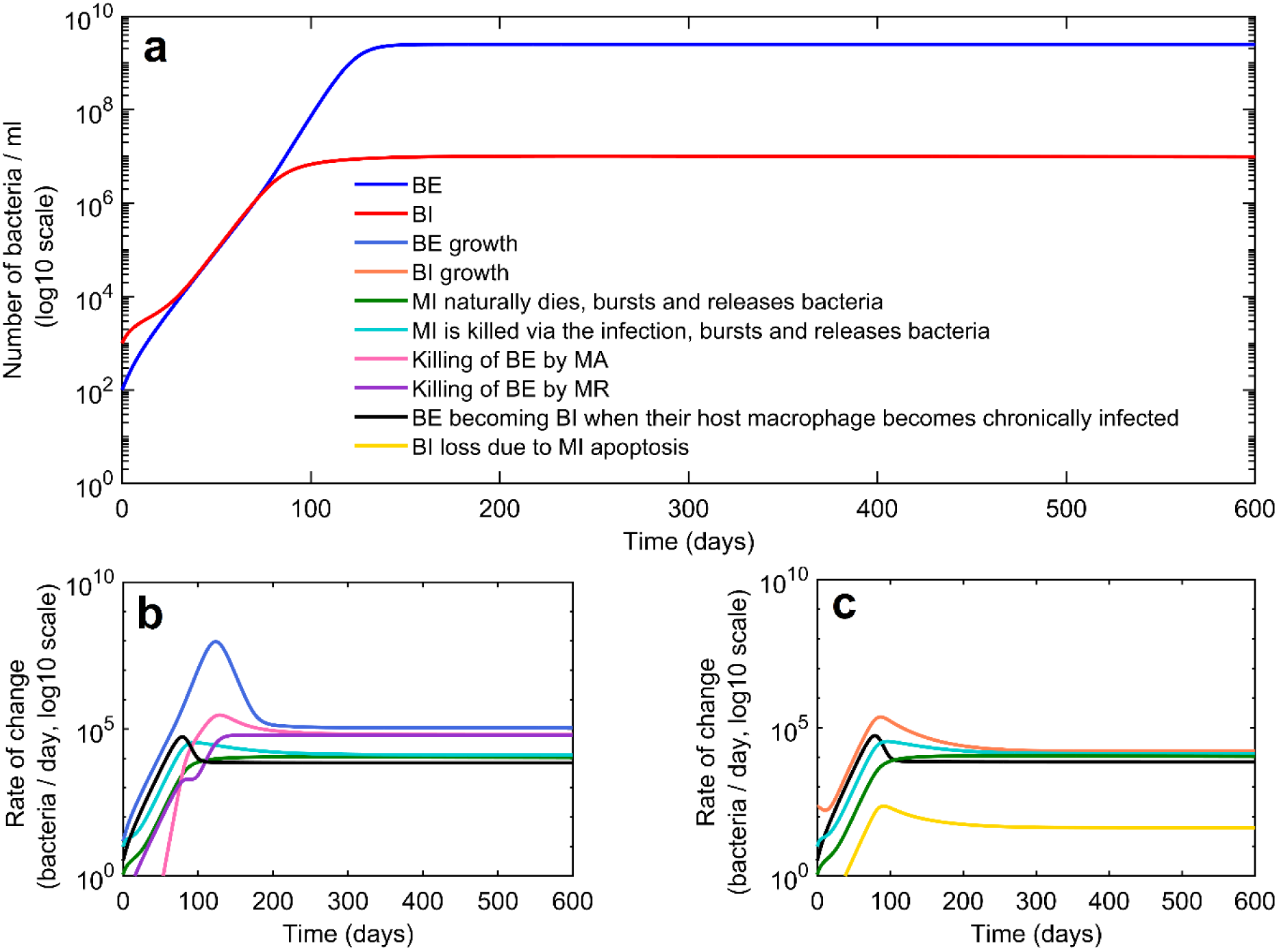
Panel (a) shows the dynamics of extracellular and intracellular *Mtb* during active tuberculosis disease, with extracellular *Mtb* representing the majority of the bacteria population. The dynamics of BE was primarily driven by the growth rate and the killing by macrophages (panel b). In contrast, the evolution of BI was mainly governed by the growth rate and the conversion between BE and BI (e.g. invasion of BE into MR and BI released from MI due to MI rupture either naturally or via TB infection) (panel c). BE, extracellular bacteria; BI, intracellular bacteria; MA, activated macrophage; MI, chronically infected macrophage; MR, resting macrophage; *Mtb*, *Mycobacterium tuberculosis*.

After between-subject variability was introduced in the model, we simulated the dynamics of *Mtb* in 500 hypothetical patients (Fig. 3). Although substantial variations in the exponential growth phase of BE were observed, the between-subject variability reduced dramatically when BE became stable (Fig. 3), indicating that the long-term dynamics are relatively more predictable than short-term transients. The correlations between variability in immune response input parameters and *Mtb* dynamics are weak (Section 5 of the Supplemental Material).

**FIG 3.**
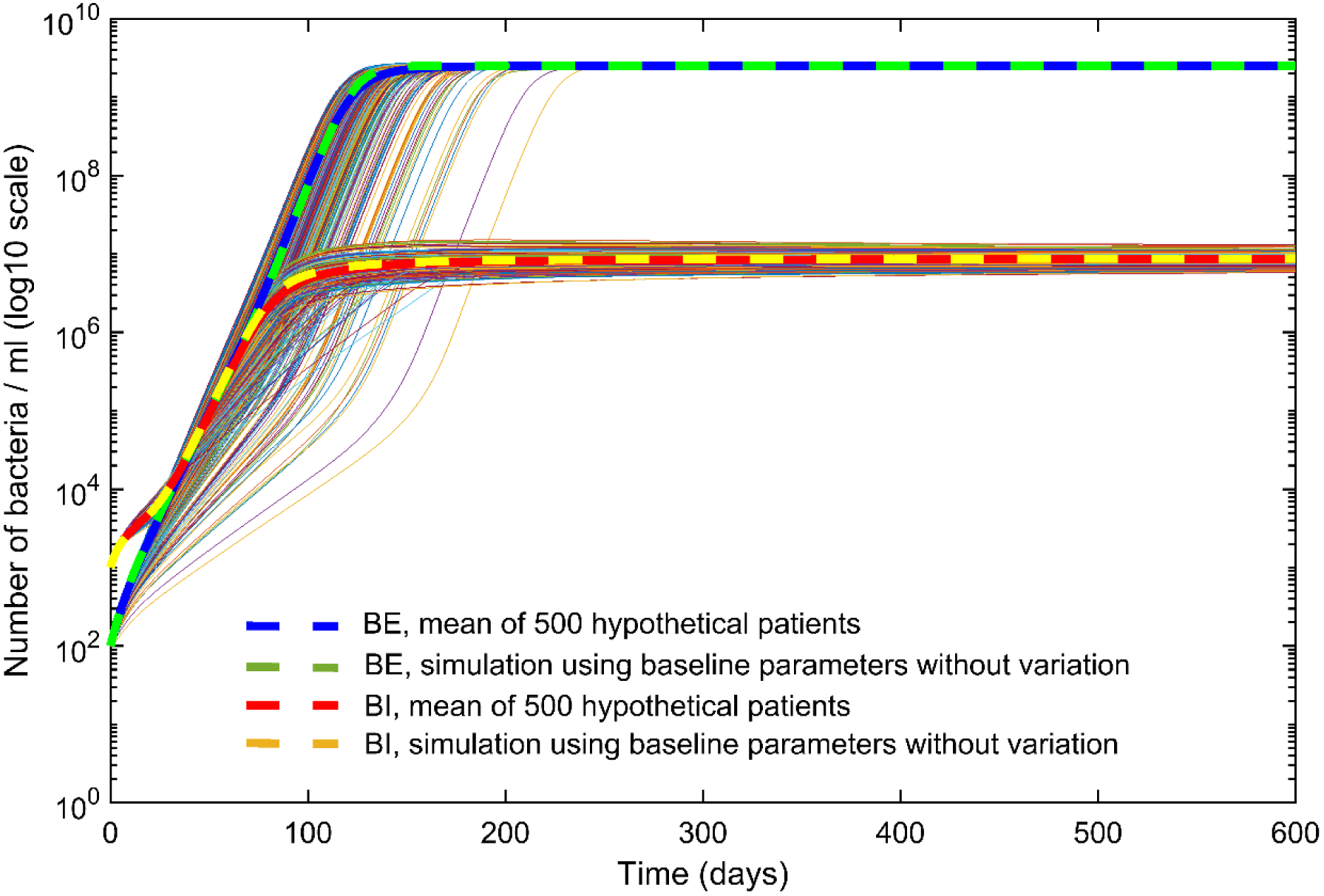
Comparison of model outputs for bacterial dynamics with and without between-subject variability. BE, extracellular bacteria; BI, intracellular bacteria.

### Simulations with treatment

Fig. 4 shows excellent agreement in treatment success rates between our model and those reported in the clinical trial by the Damien Foundation in Bangladesh (6). Fig. 5 shows the dynamics of *Mtb* when patients are treated with the nine-month Bangladesh regimen. Eighty-six percent of the patients achieved complete bacterial clearance (treatment success rate = 86%) and the median time to bacterial clearance was 69 (95% confidence interval 41-113) days. In order to test whether the results were representative, we repeated the simulation 10 times, each of which was based on a new set of 500 hypothetical patients with parameters drawn from the same parameter space using Latin Hypercube Sampling. We found that both the success rate of 86% and median time to bacterial clearance of 69 days were consistent with the results of those additional simulations (treatment success rates ranged from 84.4% to 86.4% and median time to bacterial clearance ranged from 63 to 69 days). Of note, the results are also highly consistent with clinical trials and observations from real-life clinical practice, which report treatment success rates of this regimen ranging from 84% to 88% and a median time to sputum culture conversion ranging from 30 to 152 days (5-7, 34, 35). The majority (421/430) of patients with treatment success achieved bacterial clearance during the intensive phase of treatment. The remaining nine cases did so early in the continuation phase.

**FIG 4.**
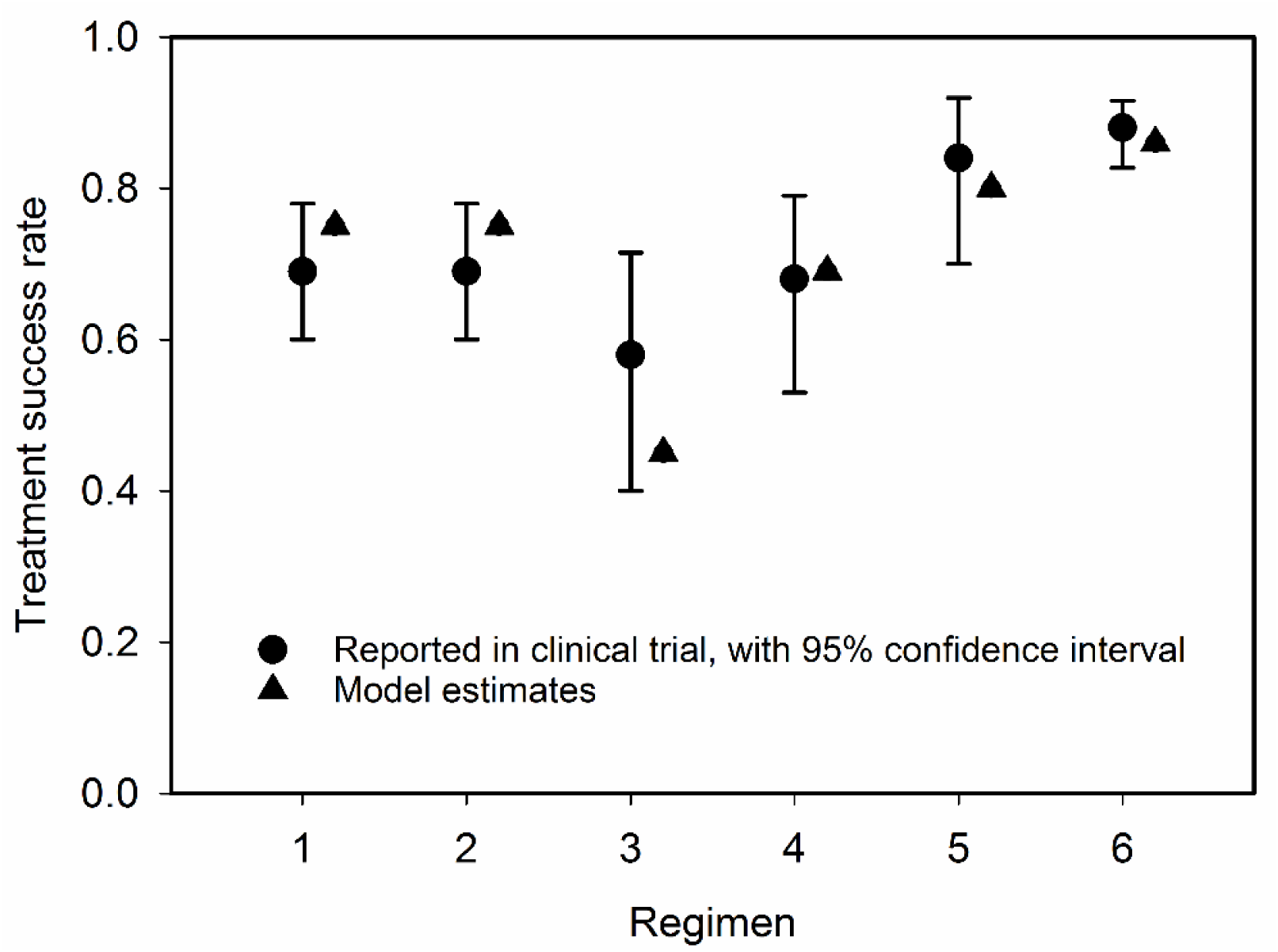
Comparison of treatment success rates between those reported in the clinical trial by the Damien Foundation in Bangladesh (6) and model estimates. Refer to Table 1 for description of the regimens.

**FIG 5.**
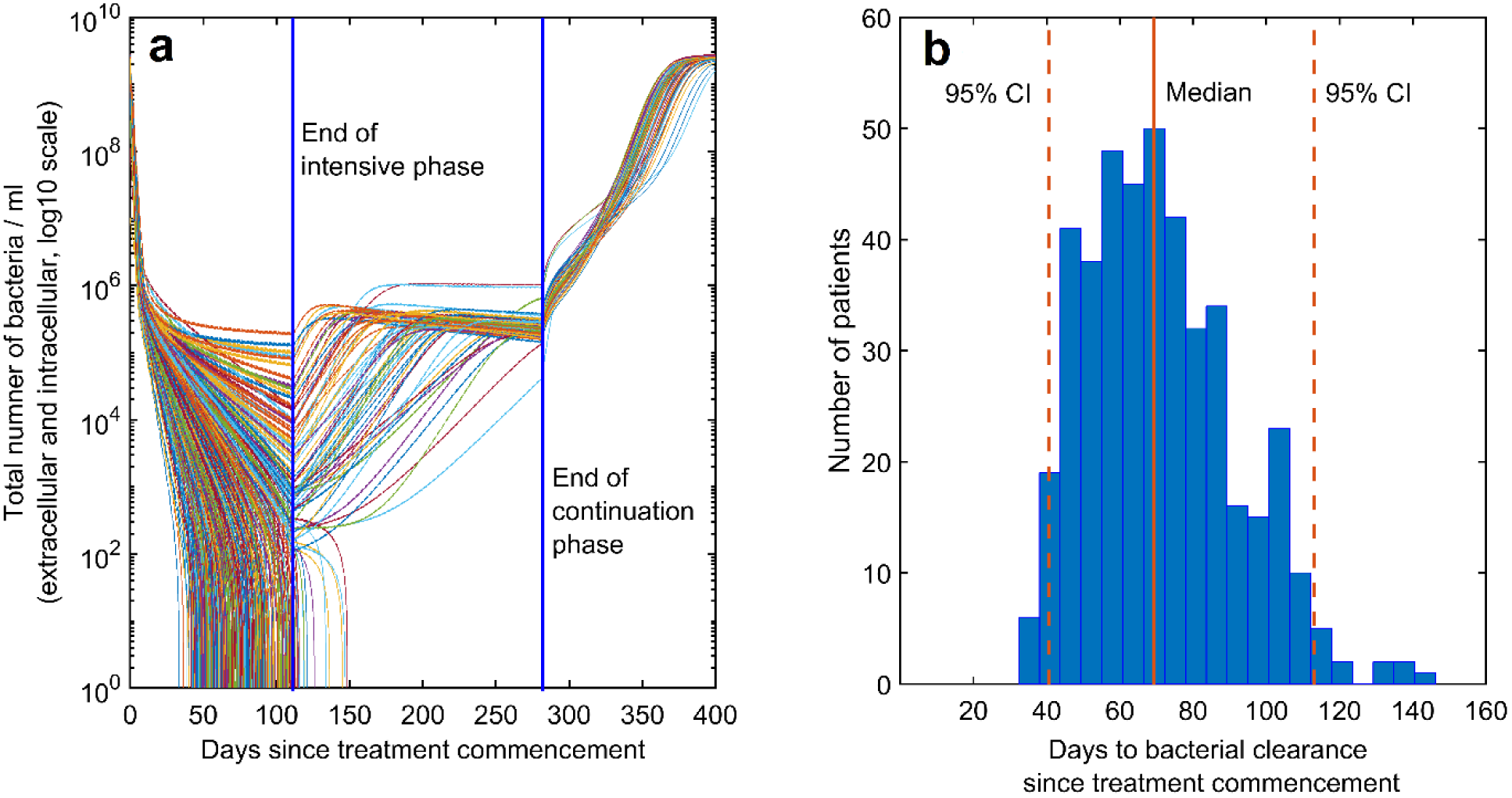
(a) The dynamics of *Mtb* (total bacteria load, including both intracellular and extracellular bacteria) in the presence of the nine-month regimen. (b) Distribution of time to bacterial clearance among the 500 simulated individuals. CI, confidence interval; *Mtb*, *Mycobacterium tuberculosis*.

An in-depth analysis of patients with treatment failure revealed that these patients achieved a lower cumulative drug-mediated killing rate than those with treatment success in both the intensive and continuation phases of treatment, with the biggest difference in killing rates between the two groups observed in the intensive phase (Fig. 6). Intermittent dosing (three times weekly or weekly) of MXF during the continuation phase did not compromise treatment efficacy compared to standard daily dosing of this drug (Fig. 7). Omitting INH markedly reduced the treatment success rate to just 34% and prolonged the median time to bacterial clearance to 92 days.

**FIG 6.**
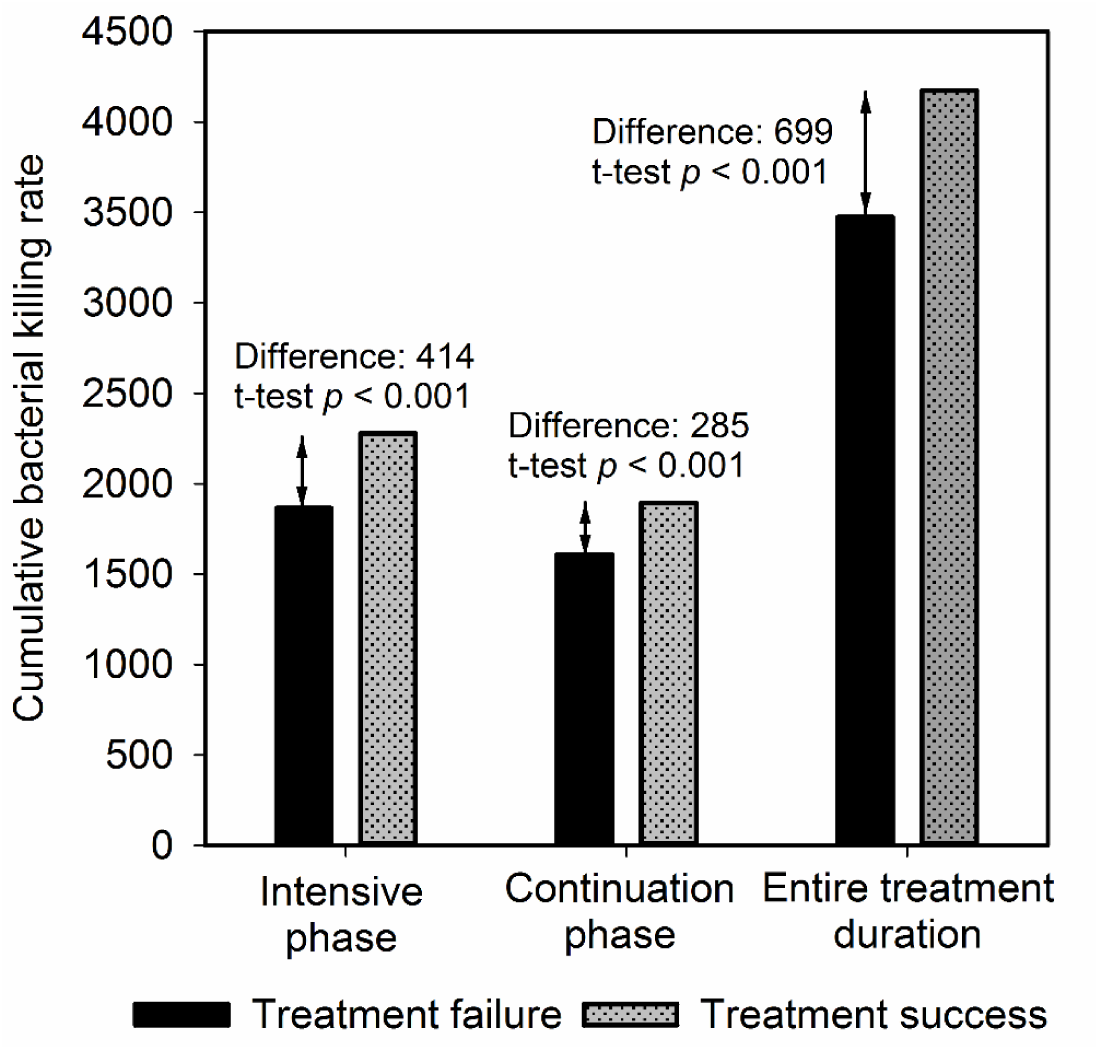
Comparison of cumulative drug-mediated bacterial killing rates between treatment failure and treatment success cases of the nine-month Bangladesh regimen. Each bar represents the mean of the cumulative bacterial killing rate of all patients in the respective treatment outcome groups (success or failure).

**FIG 7.**
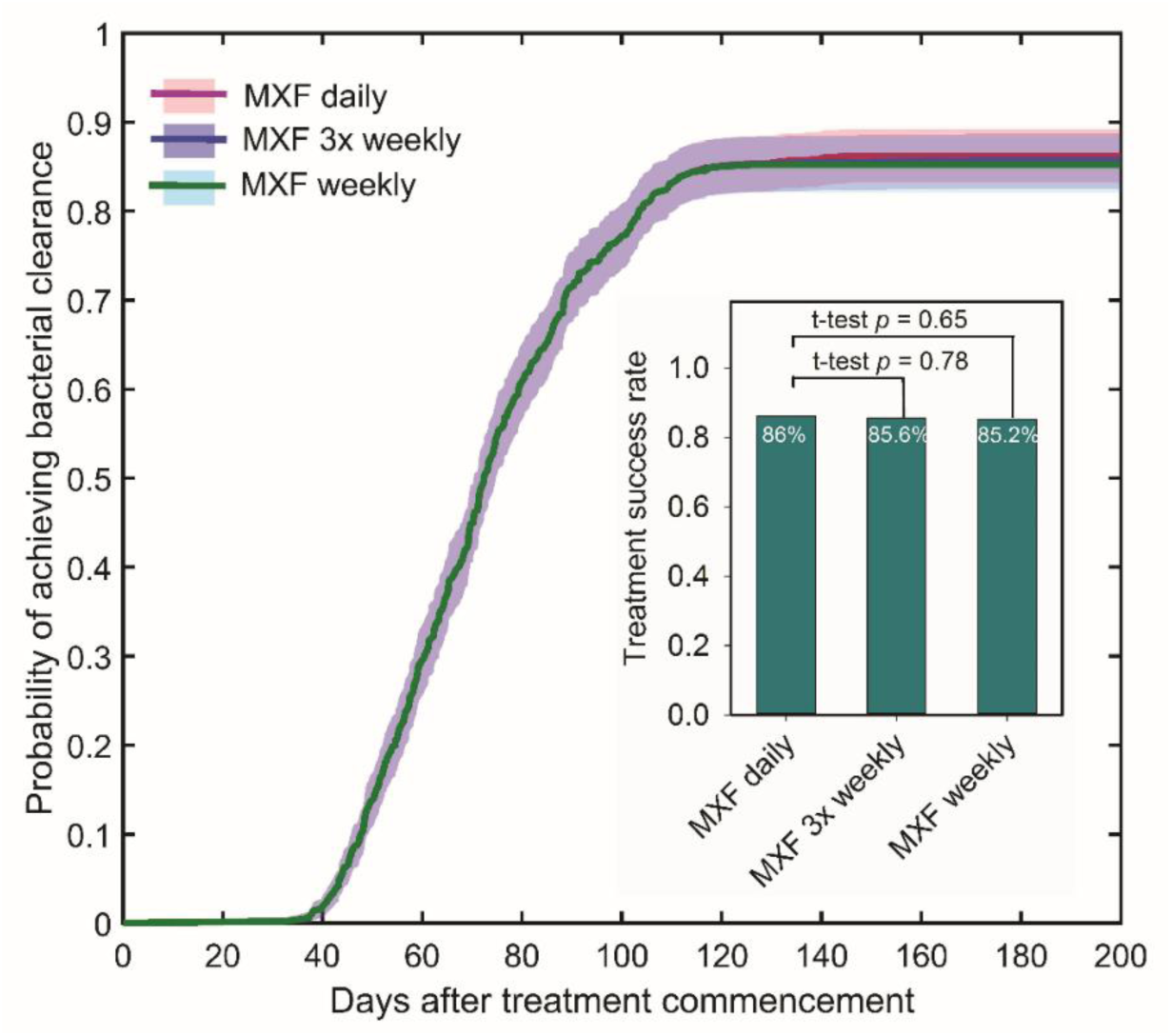
Comparisons of time to bacterial clearance and treatment success rates between daily, three times weekly (3x weekly) and weekly administrations of moxifloxacin (MXF) in the nine-month regimen. Shaded areas represent the 95% confidence intervals. Note that the curves of MXF daily, three times weekly and weekly are almost overlapping.

Table 2 shows the outcomes of treatment with the proposed short-course regimens that contain BDQ. All patients who received regimens N1-N3 achieved successful treatment (treatment success rate = 100%). In these regimens, BDQ was given daily for at least the first two weeks of treatment, and the total treatment durations ranged from 26 to 22 weeks. Treatment duration can be reduced further to just 18 weeks (approximately 4 months) while still maintaining a very high treatment success rate (100% for daily BDQ for two weeks during the intensive phase in regimen N4, and 95% when BDQ was given daily for one week during the intensive phase in regimen N5). Most of the killing effect of the investigated BDQ-containing regimens is attributable to PZA and BDQ, followed by KNM, INH and MXF; whereas the relative contribution of CFZ to the total killing effect of the regimens is negligible (Fig. 8).

**TABLE 2.** Treatment outcomes with various short-course BDQ-containing regimens

| Regimen | Intensive phase<sup>*;§</sup> | Continuation phase<sup>†;§</sup> | Treatment success rate | Median time to clearance (days) |
| --- | --- | --- | --- | --- |
| N1 | 8 MXF, CFZ, PZA, INH, KNM<br>2 BDQ (daily) | 18 MXF, CFZ, PZA<br>18 BDQ (3x weekly) | 100% | 23 |
| N2 | 8 MXF, CFZ, PZA, INH, KNM<br>2 BDQ (daily) | 18 MXF, CFZ, PZA | 100% | 24 |
| N3 | 4 MXF, CFZ, PZA, INH, KNM<br>2 BDQ (daily) | 18 MXF, CFZ, PZA | 100% | 23 |
| N4 | 2 MXF, CFZ, PZA, INH, KNM<br>2 BDQ (daily) | 16 MXF, CFZ, PZA | 100% | 27 |
| N5 | 2 MXF, CFZ, PZA, INH, KNM<br>1 BDQ (daily) | 16 MXF, CFZ, PZA | 95% | 33 |
| N6 | 2 MXF, CFZ, PZA, INH, KNM<br>1 BDQ (3x weekly) | 16 MXF, CFZ, PZA | 69% | 44 |

\*All drugs were given daily, except for BDQ of which dosing schedule was indicated in brackets.
§Numbers in front of regimen indicate weeks. †All drugs were given daily, except for MXF which was given once weekly and BDQ of which dosing schedule was indicated in brackets. BDQ, bedaquiline; CFZ, clofazimine; INH, isoniazid; KNM, kanamycin; MXF, moxifloxacin; PZA, pyrazinamide; 3x weekly, three times weekly.

**FIG 8.**
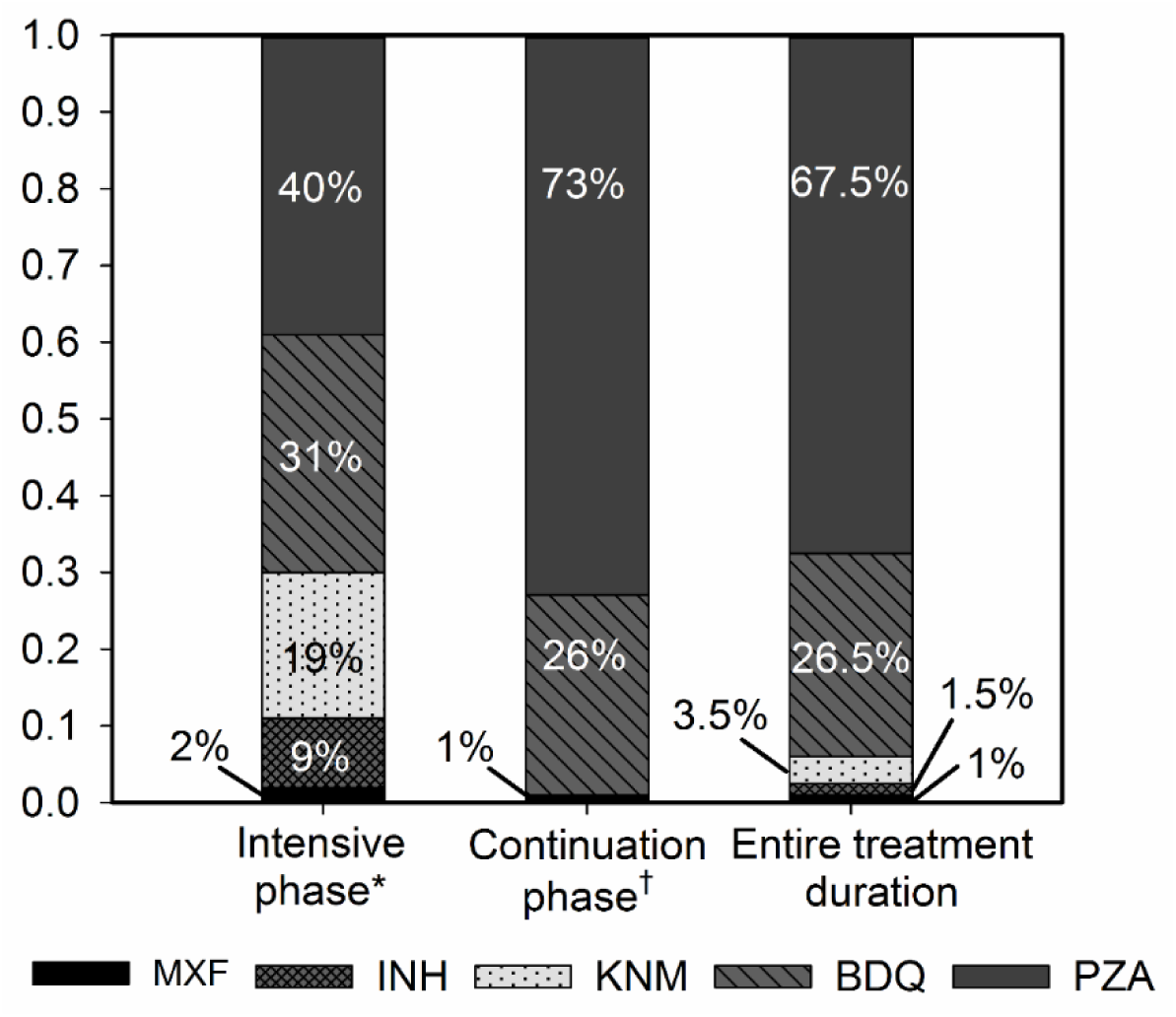
Relative contribution of individual drugs to regimen’s mycobacterial killing effect. The relative contribution of CFZ to the total killing effect is negligible and therefore is not shown. Data from regimen N5 were used. *Intensive phase includes two weeks of daily MXF, CFZ, PZA, INH, KNM and one week of daily BDQ. †Continuation phase includes 16 weeks of daily CFZ, PZA and three times weekly MXF. BDQ, bedaquiline; CFZ, clofazimine; EMB, ethambutol; INH, isoniazid; KNM, kanamycin; MXF, moxifloxacin; PZA, pyrazinamide.

## DISCUSSION

This study is the first to use a mathematical modelling approach incorporating the intrahost immune response, PK and PD to investigate the effect of short-course regimens containing the new anti-TB drug BDQ, and its potential for further shortening (to less than nine months) treatment for MDR-TB. Our simulations suggest that BDQ can help reduce treatment duration to four months while improving treatment success rate to 95% (when BDQ is given daily for one week during intensive therapy) or 100% (when BDQ is given daily for two weeks during intensive therapy) compared to the reported 84-88% success rate of the nine-month Bangladesh regimen (5-7). These results, by establishing likely efficacy and alleviating any potential ethical concerns regarding clinical trials, provide a rationale in support of future clinical investigations of short-course regimens containing BDQ for MDR-TB. Four-month treatment of MDR-TB would greatly simplify treatment supervision, improve treatment compliance and patient’s comfort, save costs, and allow more free time and earning potential for patients and caretakers. The four-month regimen would also be expected to minimise acquisition of additional resistance (36). Our model also predicts that dosing frequency of MXF during continuation therapy could be reduced from the standard daily to once weekly administration while maintaining efficacy of treatment. Intermittent dosing of MXF would result in cost savings due to a reduction in the time and resources consumed with daily administration of the drug. Less frequent dosing of MXF would also likely further improve patient adherence (37). Information on the cardiac safety, especially QTc prolongation, of BDQ in combination with CFZ or fluoroquinolones (e.g. MXF) remains scarce. However, available data suggest that such adverse events are rare and not attributable to the administration of these drugs (9-11, 38-40). Nevertheless, electrocardiography monitoring should be performed during therapy with these drugs, especially in patients with predisposing factors for QTc prolongation.

Our model contains more than one hundred parameters, making it computational intractable to conduct an exhaustive uncertainty analysis. In order to adequately quantify uncertainty, we used various resources to determine the parameter values and their plausible ranges. For the parameters related to the bacterial dynamics and the immune response, most parameters were adapted from Wigginton and Kirschner (15) where the authors provided detailed estimations and justifications for the parameters. Most of the PK and PD parameters were directly drawn from published experimental studies with the exception that the PD parameters for BDQ, CFZ, KNM and EMB were not immediately available and needed to be estimated based on *in vitro* data (see Section 6 of the Supplemental Material for details). We did not calibrate our model to any pre-defined outcome values. As such, our application of the model to simulate all the regimens investigated in the landmark clinical trial by Damien Foundation in Bangladesh (6) may be considered as an independent test of its validity. We found that the predicted treatment outcomes were highly consistent with previous clinical findings, constituting a successful test of the model. This demonstrates the reliability of the model to capture the dynamics of natural TB infection and make predictions.

As with all models (mathematical or otherwise), our mathematical model is a deliberate simplification of the true biological system, developed to aid understanding and guide experiment. In that context, we note that certain aspects of the immune response to TB remain poorly understood (41). We have focussed on the role of macrophages, T-cells and four cytokines. We did not include every type of cell and cytokine that may potentially play a role in the body’s defence mechanism against *Mtb* infection such as neutrophils, natural killer cells, IL-2 and TGF-*β* (15). Nevertheless, our model represents the most comprehensive model of the immune response to TB to date. It can be readily adapted to incorporate other cells when new data become available. We did not consider the effect that potential interactions between the included drugs may have on their overall anti-mycobacterial activity; nor did we account for the acquisition of additional drug resistance during treatment. These are important areas for future research.

## ACKNOWLEDGEMENTS

We gratefully acknowledge the financial support provided by the PRISM^2^ Seed Funding Grant from the Australian Government National Health and Medical Research Council Centre of Research Excellence in Infectious Diseases Modelling to Inform Public Health Policy (granted to TND and PC), and the Early Career Researcher Grant from the University of Melbourne (granted to TND). All authors declare that they have no competing interests.

